# Eye-position error influence over “open-loop” smooth pursuit initiation

**DOI:** 10.1101/404491

**Authors:** Antimo Buonocore, Julianne Skinner, Ziad M. Hafed

## Abstract

The oculomotor system integrates a variety of visual signals into appropriate motor plans, but such integration can have widely varying time scales. For example, smooth pursuit eye movements to follow a moving target are slower and longer-lasting than saccadic eye movements, and it has been suggested that initiating a smooth pursuit eye movement involves an obligatory open-loop interval, in which new visual motion signals presumably cannot influence the ensuing motor plan for up to 100 ms after movement initiation. However, this view runs directly contrary to the idea that the oculomotor periphery has privileged access to short-latency visual signals. Here we show that smooth pursuit initiation is sensitive to visual inputs, even in “open-loop” intervals. We instructed male rhesus macaque monkeys to initiate saccade-free smooth pursuit eye movements, and we injected a transient, instantaneous eye position error signal at different times relative to movement initiation. We found robust short-latency modulations in eye velocity and acceleration, starting only ∼50 ms after transient signal occurrence, and even during “open-loop” pursuit initiation. Critically, the spatial direction of the injected position error signal had predictable effects on smooth pursuit initiation, with forward errors increasing eye acceleration and backwards errors reducing it. Catch-up saccade frequencies and amplitudes were also similarly altered ∼50 ms after transient signals, much like well-known effects on microsaccades during fixation. Our results demonstrate that smooth pursuit initiation is highly sensitive to visual signals, and that catch-up saccade generation is reset after a visual transient.

## Introduction

An intriguing property of motor circuits is that they have privileged access to sensory signals. For example, neck muscles and related upper body motor synergies, which re-orient the upper body towards behaviorally-relevant stimuli, exhibit early stimulus-locked responses after visual flashes (Corneil, 2004; Corneil et al., 2013; Corneil and Munoz, 2014). These responses, arriving less than 100 ms after stimulus onset, occur almost as early as neural responses in primary visual areas (Corneil, 2004; Boehnke and Munoz, 2008; Corneil and Munoz, 2014). Even within the realm of visually-guided oculomotor behavior, our topic of interest here, smooth ocular following can be elicited by full-field image motions within the same time frame (Kawano and Miles, 1986), and much shorter response latencies can be expected if other sensory modalities were to influence eye movements (as in the case of vestibular ocular reflexes; Lisberger, 1984; Cullen et al., 1991; Snyder and King, 1992; Minor et al., 1999). All of these observations are consistent with the fact that brainstem oculomotor control circuitry are riddled with eye-movement related neurons exhibiting both visual and eye-movement related discharge in the very same neurons. In fact, such concomitance of visual and motor signals in the same neurons can occur even in pre-motor nuclei (Evinger and Kaneko, 1982; Everling et al., 1998; Missal and Keller, 2002), a mere single synapse away from motor neurons innervating the ocular muscles.

Intuitively, access to the sensory periphery by motor systems makes sense. In the case of vision and the oculomotor system, such access allows rapid resetting of oculomotor plans in order to make appropriate orienting reflexes (Reingold and Stampe, 1999, 2002; Buonocore and McIntosh, 2008; Rolfs et al., 2008; Hafed and Ignashchenkova, 2013). This resetting can even alter the kinematics of already-programmed saccades (Edelman and Xu, 2009; Guillaume, 2012; Buonocore et al., 2016; Buonocore et al., 2017b) well before most higher-level cortical visual areas finalize processing of the appearing stimuli (Schmolesky et al., 1998). That is, within a mere ∼50-100 ms after stimulus onset, a full processing cycle from visual input all the way to eye movement output takes place (Fischer and Boch, 1983; Boch et al., 1984; Fischer and Ramsperger, 1984; Tian et al., 2018). This rapid access to visual inputs imparts on the oculomotor system a highly efficient pathway for behavioral re-orienting (Buonocore et al., 2017a).

Despite the above, and even though recent research has shown that smooth pursuit eye movements do share neural resources with saccades (Krauzlis, 2004), the different time scales between these two types of eye movements (brief and ballistic for saccades, versus long-lasting and slow for smooth pursuit) have historically led to descriptions of certain categorical differences between these two types of eye movements. Namely, early pioneering work on smooth pursuit initiation (Rashbass, 1961; Lisberger and Westbrook, 1985; Tychsen and Lisberger, 1986) has overwhelmingly become interpreted as suggesting that such initiation possesses an obligatory “open-loop” interval of ∼100 ms (in monkeys; ∼130 ms in humans), before which new visual motion signals cannot influence the motor behavior. While contradicting the idea that the oculomotor periphery has privileged access to visual signals, this concept has become pervasive enough that a number of studies have explicitly relied on it to probe perception by means of visual stimulation, and assuming that smooth pursuit initiation would be unaffected (e.g. Spering et al., 2006; Schütz et al., 2007; Schutz et al., 2008; Braun et al., 2017).

Our goal here was to explicitly demonstrate that smooth pursuit initiation is indeed sensitive, and with short latencies, to visual stimuli. We introduced instantaneous visually-defined eye position errors during pursuit initiation. We found a systematic directional influence of such signals, and with latencies of only 50-80 ms. These results, directly paralleling saccadic ones (Reingold and Stampe, 2002; Buonocore and McIntosh, 2008; Edelman and Xu, 2009; Bompas and Sumner, 2011; Buonocore et al., 2017b), suggest that the concept of “open-loop pursuit” may be too coarse to accurately describe the mechanisms of smooth pursuit initiation.

## Materials and Methods

### Animal preparation and laboratory setup

We recorded eye movements with high spatial and temporal precision from two male rhesus macaque monkeys (monkey A and monkey M) aged 7 years. The monkeys were implanted, under general anesthesia and using sterile surgical techniques, with a titanium head-post attached to the skull with titanium skull screws (all implanted under the skin and muscle surrounding the skull). The head-post allowed stabilizing the animals’ head position during the experiments. In a subsequent surgery, one eye in each animal was implanted with a scleral search coil to allow tracking eye movements using the magnetic induction technique (Fuchs and Robinson, 1966; Judge et al., 1980). The animals are part of a larger neurophysiology experiment for which the behavioral training needed for the current study was a necessary prerequisite. The experiments were approved by ethics committees at the regional governmental offices of the city of Tuebingen, and these experiments were in accordance with European Union guidelines on animal research, as well as the associated implementations of these guidelines in German law.

During the experiments, the animals were seated in a primate chair 74 cm from a CRT computer monitor in an otherwise dark room. The monitor had a pixel resolution of 34 pixels/deg and a refresh rate of 120 Hz, and stimuli were presented over a uniform gray background (29.7 Cd/m^2^). A small white spot (5 x 5 min arc square) having 86 Cd/m^2^ luminance was used as either a fixation spot or the moving target for smooth pursuit eye movements (see *Behavioral tasks* below). Other stimuli that we used consisted of white flashes of different sizes and positions, as described in *Behavioral tasks* below; they all had the same luminance as the spot (86 Cd/m^2^). The CRT monitor viewed by the monkeys was controlled by a computer running Matlab’s Psychophysics Toolbox (Brainard, 1997; Pelli, 1997; Kleiner et al., 2007). This computer was in turn receiving commands from a real-time I/O system from National Instruments (Austin, USA), which was maintaining control of the display on a frame-by-frame basis. The system was programmed in-house and described recently (Chen and Hafed, 2013; Tian et al., 2016).

### Behavioral tasks

#### Experiment 1: Saccade-free smooth pursuit initiation with transient position error signals

We employed a paradigm eliciting smooth (i.e. saccade-free) ocular following of a moving target while we introduced a transient eye position error signal in a specific direction (Fig. 1A). Trials started with a white fixation spot presented at the center of the display over a uniform gray background. After the monkey aligned gaze with the fixation spot by 700-1000 ms, the spot jumped by 4 deg either to the right or left of fixation and simultaneously started moving horizontally (at a speed of 20.4 deg/s) in the opposite direction. The step size was chosen such that by the time a smooth pursuit eye movement (in the target motion direction) was initiated, no initial catch-up saccade was needed (Rashbass, 1961). This allowed us to achieve saccade-free smooth pursuit initiation velocities and, therefore, to explore how instantaneous eye position error signals could modify such smooth velocity initiation.

**Figure 1.**
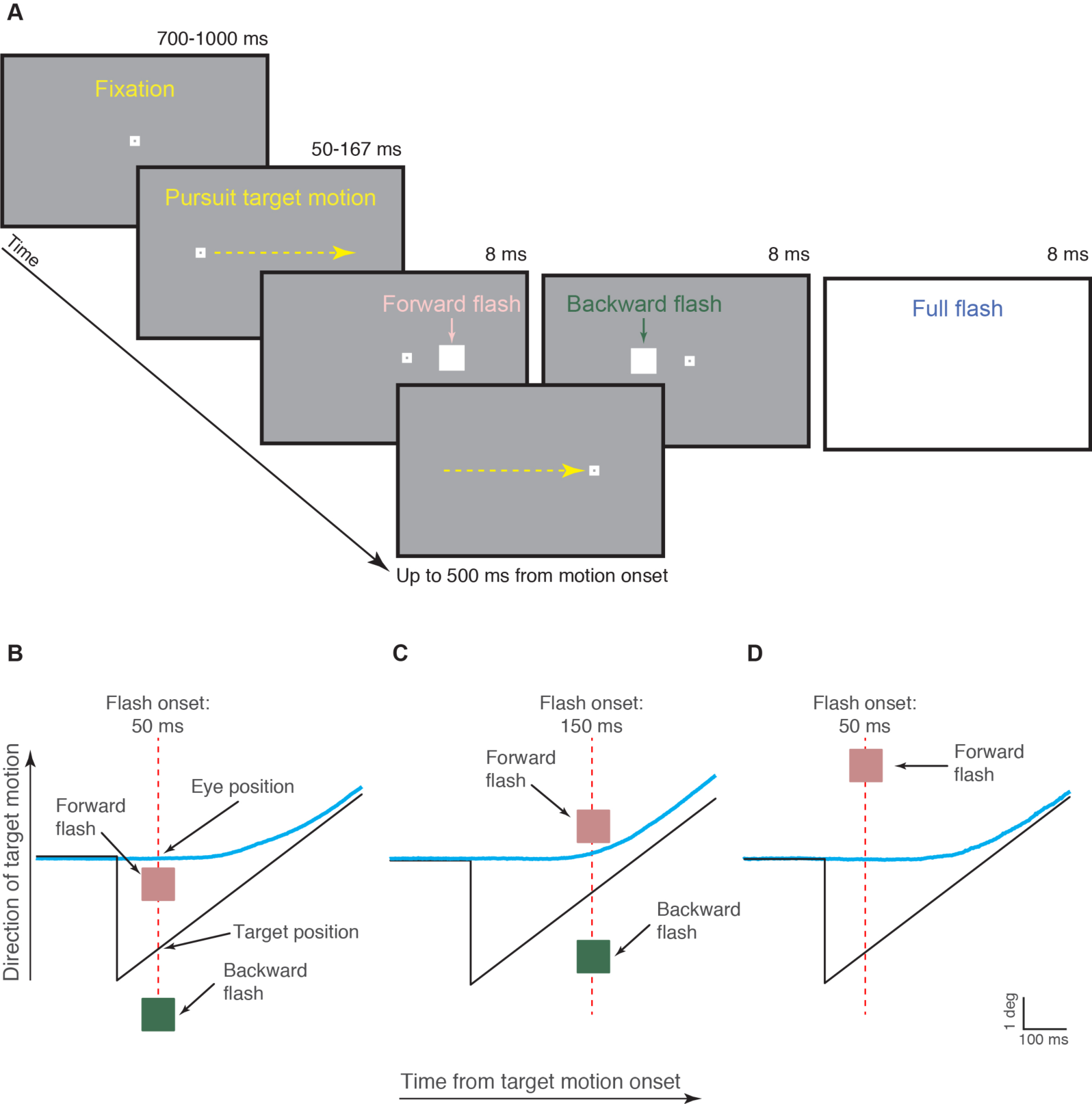
Methods. (A) Each trial in Experiment 1 started with the monkey fixating display center for 700-1000 ms. The fixation spot then jumped right or left and instantaneously started moving in the opposite direction. At a random time 50-167 ms after motion onset, a 1 × 1 deg flash was presented for 8 ms at 2.1 deg either in front of (pink label) or behind (green label) the target. In some trials, the flash covered the entire screen (light blue label). (B-D) Depending on instantaneous gaze position at the time of flash onset, “forward” and “backward” flashes in the task were either in front of or behind gaze. (B) Example eye trace (blue) relative to target position (black) for a flash presented at 50 ms after motion onset. In this case, both forward and backward flashes were spatially located behind instantaneous eye position. (C) Example eye trace for a flash presented at 150 ms after motion onset. The forward flash was now in front of instantaneous eye position while the backward flash was behind it. (D) Example eye trace for a flash presented at 50 ms after motion onset and a forward flash being located at 6 deg in front of target position. In this case, the forward flash was in front of eye position even for this early flash time.

Instantaneous eye position error signals were introduced by interleaving trials in which the target motion above was accompanied, for a single display frame (∼8 ms), by a white flash of the same luminance as the pursuit target spot (Fig. 1A). The flash, appearing 50-167 ms (in steps of ∼8 ms) after target motion onset, could be a full-screen flash or a square of 1 x 1 deg dimensions centered on a position either 2.1 deg in front of or 2.1 deg behind the instantaneous pursuit target position (along the motion trajectory, as in Fig. 1B, C; in monkey M, we also ran the experiment with the square appearing at 6 deg, instead of 2.1 deg, in front of or behind the instantaneous pursuit target position, as in Fig. 1D). Thus, depending on instantaneous gaze position at the time of flash onset, the “forward” and “backward” flashes allowed us to map different retinotopic positions of visual transients during smooth pursuit initiation (Fig. 1B-D).

The monkeys were rewarded for maintaining their gaze within 3 deg from the pursuit target position until trial end (occurring 500 ms from target motion onset). This virtual gaze window for monitoring monkey behavior was relatively relaxed because we found that the flash could momentarily “attract” gaze position (with both smooth velocity and saccadic effects; see Results), and also because the monkeys were highly trained on accurate gaze alignment on the pursuit target anyway. Across sessions, we collected 3624 trials from monkey M and 1996 trials from monkey A in this task (with the forward and backward flashes being 2.1 deg in front of or behind the pursuit target; Fig. 1B, C). We also collected 3705 trials from monkey M in the variant with forward and backward flashes being 6 deg in front of or behind the pursuit target (Fig. 1D). The four experimental conditions (normal saccade-free pursuit initiation without a flash, initiation with a full-screen flash, initiation with a forward flash, or initiation with a backward flash) were randomly interleaved within a session. The monkeys typically completed ∼300 trials of this task per session, with the remainder of the session dedicated to the other tasks below, or to other neurophysiology-related tasks not described in this study.

#### Experiment 2: Saccadic smooth pursuit initiation with transient position error signals

We repeated the same experiment, except that there was no initial step in target position before the target motion onset. That is, in the present experiment, the monkeys steadily fixated the fixation spot at the center of the display, and the spot then started moving at a constant speed directly from the center. Because there is an unavoidable latency in reacting to the spot’s motion onset, the monkeys had to perform an initial catch-up saccade as part of initiating their smooth tracking behavior, and our purpose was to explore the influence of instantaneous eye position error signals introduced by flashes on such an initial catch-up saccade. All other details were identical to those described above for Experiment 1. We collected 1824 trials from monkey M and 1511 trials from monkey A in this task, and the forward and backward flashes in this experiment were only at a 2.1 deg eccentricity relative to the pursuit target.

#### Experiments 3: Steady-state pursuit accompanied by a transient flash stimulus

To compare the effects of transient instantaneous eye position error signals on smooth pursuit initiation to effects during steady-state pursuit, we ran an additional version of the above experiments. In Experiment 3, the flashes could appear during steady-state smooth pursuit as opposed to during smooth pursuit initiation. Trials started with the fixation spot presented at +/-7 deg horizontally from display center, to allow for a long period of sustained smooth pursuit spanning large portion of the horizontal extent of our display area. After 300-500 ms of fixation, the target started moving towards (and past) display center at a speed of 12 deg/s. Note that this speed was slower than that in Experiments 1 and 2 because we wanted to establish a sufficiently long duration of sustained steady-state smooth pursuit within trials. The motion lasted for ∼1500-1800 ms, and at a random time during steady-state smooth pursuit (up to 1500 ms from motion onset; in steps of ∼8 ms), a flash could appear as in Experiments 1 and 2 above (forward and backward flashes were at 2.1 deg). We explored the effects of the flash and its type (forward, backward, or full-screen) on both smooth velocity effects as well as on catch-up saccades. We collected 5085 trials from monkey M and 2849 trials from monkey A in this experiment.

### Data analysis

#### Detecting catch-up saccades and microsaccades

We numerically differentiated our digitized eye position signals to obtained estimates of eye velocity and acceleration. We used velocity and acceleration criteria to detect microsaccades and catch-up saccades, as described previously (Krauzlis and Miles, 1996a; Hafed et al., 2009; Chen and Hafed, 2013). We then manually inspected all data to correct for misses or false detections.

#### Measuring smooth eye velocity effects

In Experiment 1, our aim was to isolate the influence of transient instantaneous eye position error signals on smooth pursuit initiation. Our task design minimized the occurrence of catch-up saccades. In the analysis, we replaced all saccades and catch-up saccades from the eye trace with Not-a-Number (NaN) labels that extended in duration to +/-50 ms around each detected saccade. This way, we made sure that smooth pursuit eye velocity averages were not influenced by high velocity changes due to the rare saccades that may have occurred. This procedure avoided catch-up saccades during initiation, and, equally important, our exclusion of microsaccades also avoided the potential influence of these fixational eye movements (occurring near target motion onset) on visual sensitivity and behavior (Hafed and Krauzlis, 2010; Hafed, 2013; Chen et al., 2015; Hafed et al., 2015; Tian et al., 2016; Chen and Hafed, 2017; Tian et al., 2018). We then averaged the radial eye velocity traces aligned on motion onset and/or flash onset for the different trial types (e.g. no flash, full-screen flash, forward flash, or backward flash). The no flash eye velocity traces acted as our baseline, to which we compared velocities when there was a flash during pursuit initiation.

We also repeated the same analyses above but after explicitly removing every trial (in its entirety) that contained at least one saccade in the interval from −100 ms to 270 ms from target motion onset. Our results were virtually unaltered, so we elected, for the purposes of clarity, to report figures and statistics below only for this latter approach.

To compare flash conditions, we first plotted radial eye velocity after motion onset for the no flash trials as well as for either full-screen, forward, or backward flash trials. Plotting the averages side by side (e.g. see Fig. 2A in Results) allowed us to establish that there was indeed an influence of early transient flashes on “open-loop” smooth pursuit initiation. We then plotted the difference in eye velocity between flash and no flash trials. To do this, we subtracted within each session the average radial eye velocity in no flash trials from the radial eye velocity of every flash trial of a given type (e.g. for forward flashes only); we then repeated this subtraction for all sessions from a given monkey. This allowed us to achieve a population of velocity “difference” curves isolating the influence of a given flash type (e.g. Fig. 2B in Results); we estimated variance in this population by calculating the 95% confidence interval of the difference curves across all trials in all sessions.

**Figure 2.**
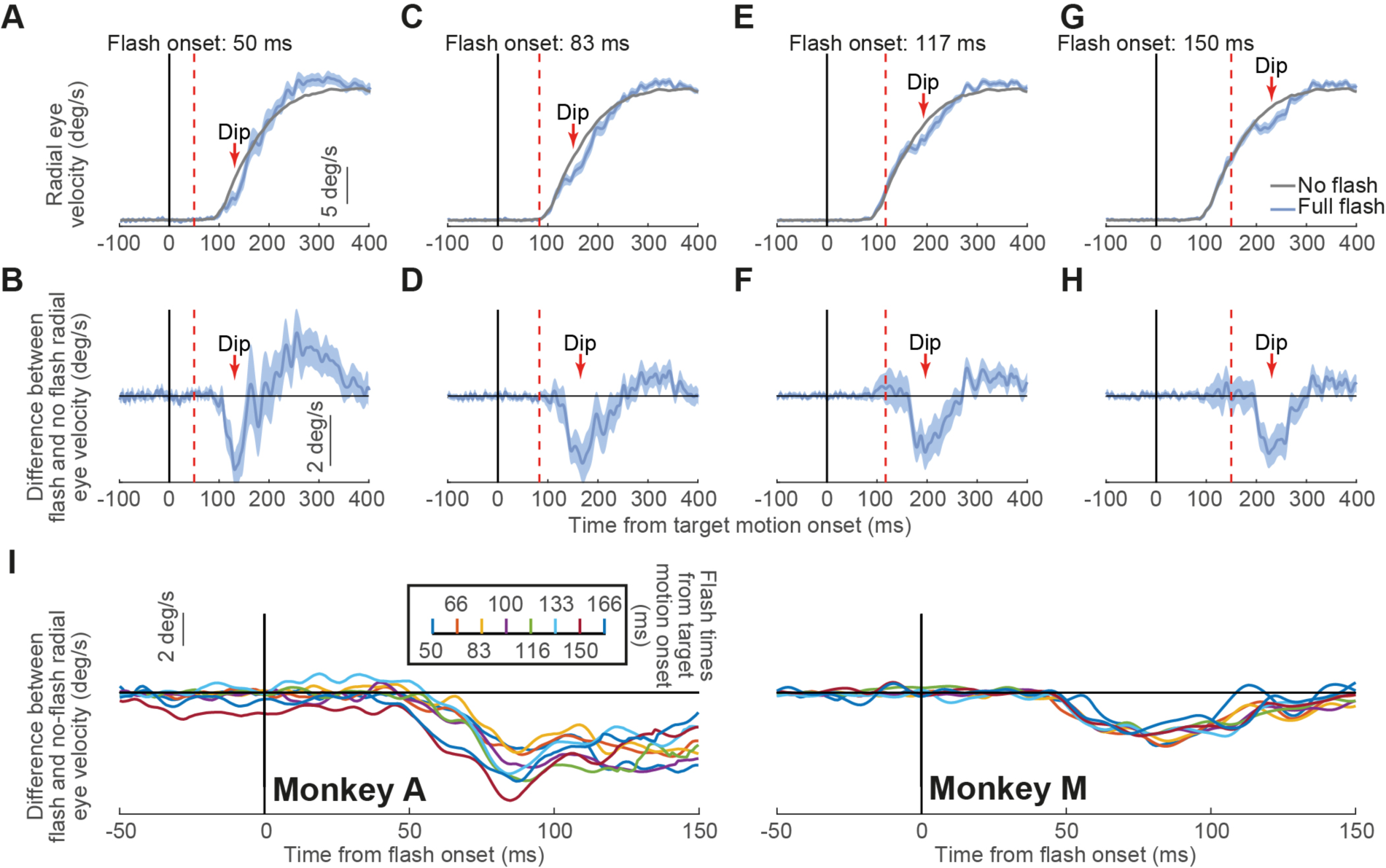
Transient disruption of open-loop smooth pursuit initiation. (A) We plotted radial eye velocity from monkey M in Experiment 1 when no flashes were presented (gray) and when full flashes were presented 50 ms after target motion onset (light blue). There was a transient dip in eye velocity relative to the gray curve starting ∼50 ms after flash onset. (B) Difference in eye velocity between flash and no-flash trials from A. The flash-induced dip in the difference curve is clearly visible. (C, D) Similar to A, B but for flashes occurring 83 ms after target motion onset. In this case, the transient disruption in open-loop pursuit initiation was slightly delayed relative to A, B because of the slight delay in flash onset time. (E, F) Same for flashes 117 ms after target motion onset. (G, H) Same for flashes after 150 ms from target motion onset. (I) In all cases, and for both monkeys, the disruption was always aligned on flash onset. We plotted eye velocity difference curves as in B, but now aligning the data on flash onset. The different colors denote different flash onset times relative to target motion onset. Regardless of flash time, the velocity difference curves always showed a dip ∼50 ms after flash onset. A series of one sample t-tests against zero showed p<0.05 (Bonferroni corrected) for all flash times for both monkeys (with the exception of flash time 83.33 and 125 in monkey A). Error bars in all panels denote 95% confidence intervals. After saccade removal (Materials and Methods), trial numbers for the 15 flash delays in the full-flash condition were 8-28 (average: 21.5) and 22-72 (average: 49) in monkeys A and M, respectively.

To summarize the impact of a given flash type on eye velocity, we identified a measurement interval (either 50-100 ms after full flash onset or 50-80 ms after backward/forward flash onset) in the velocity difference curves described above; this interval was chosen because it was the interval during which the flash exerted the influence over eye movements that we were particularly interested in investigating (see Results). For all trials of a given type (e.g. forward flash, backward flash, or full-screen flash), we averaged the eye velocity difference in this post-flash interval, and this gave us a population of measurements across trials. We then compared populations of measurements across the different flash conditions in our trial types (e.g. forward versus backward flashes) and also for different flash times relative to smooth pursuit initiation.

To compare the influence of transient instantaneous eye position error signals on smooth pursuit initiation to their influence during steady-state smooth pursuit, we repeated the above analyses but for data containing steady-state pursuit (Experiment 3). For all flash times occurring >200 ms after target motion onset (to ensure that smooth pursuit was already ongoing) and >200 ms before trial end, we plotted radial eye velocity from −100 ms to 200 ms relative to flash onset. Importantly, we removed from the eye traces all catch-up saccades (along with flanking +/-50 ms intervals) to avoid effects of high velocity changes on our measurements. We averaged trials from each type (e.g. full-screen flash, forward flash, or backward flash) to demonstrate the influence of the flash on smooth eye velocity. We also repeated the same analysis after removing every trial containing at least one saccade in the interval from −50 ms to 150 ms from flash onset. Like in Experiment 1 above, there results were again unaltered from what we report in Results.

#### Measuring effects on catch-up saccades

We analyzed catch-up saccade times and amplitudes. Since the absolute majority of catch-up saccades occurred in the direction of ongoing smooth pursuit (e.g. 97% in monkey A and 98% in monkey M from the pursuit initiation trials in Experiment 2; and 90% in monkey A and 97% in monkey M in Experiment 3), we restricted our catch-up saccade analyses to saccades in the direction of pursuit. We are confident that we did not miss catch-up saccades opposite the direction of pursuit (for which saccade-related velocity can be missed by simple velocity thresholds) because we manually inspected all trials after running our saccade detection algorithms (see above). We summarized catch-up saccade frequency of occurrence and amplitude, with and without flashes, and as a function of forward or backward flashes.

#### Experimental design and statistical analysis

Each experiment underwent within-subject analyses, and we report summaries and statistics from each subject individually. We used two male monkeys to ensure replicability of results while being constrained by the 3R (Refine, Reduce, Replace) principles for animal research. Combined with congruence of several key ingredients of our results with several prior studies in monkeys (e.g. Krauzlis and Miles, 1996b; Hafed and Ignashchenkova, 2013; Buonocore et al., 2017b) and humans (e.g. Schutz et al., 2008; Edelman and Xu, 2009; Buonocore et al., 2016; Braun et al., 2017), we have high confidence that our results from the 2 monkeys are a general property of the smooth pursuit eye movement system.

In each experiment, we randomly interleaved the relevant conditions for statistical comparisons (e.g. no flash, full-screen flash, forward flash, and backward flash in equal proportion). We also ensured a minimum of 300 repetitions per condition for the great majority of our analyses. Such a large number of repetitions made using parametric statistical comparisons in our analyses well justified. For some analyses of catch-up saccades during a “saccadic inhibition” post-flash phase of trials (Reingold and Stampe, 1999, 2000, 2002; Buonocore and McIntosh, 2008; Hafed and Ignashchenkova, 2013; Buonocore et al., 2017b), there was (by definition of the phenomenon) a relative rarity of saccades to analyze. However, we ensured statistical robustness even for these observations by increasing the number of repetitions per condition that we collected overall (see above), such that there was still statistical confidence in the relatively fewer inhibition-phase saccades that we were analyzing (we show 95% confidence intervals for all of our analyses).

Further statistical validation was obtained by performing, for each animal, two-way ANOVAs (factor “flash time” by factor “flash direction”, with the appropriate number of factor levels depending on the specific design within the experiment) on the summary statistics described above (e.g. amplitude of saccades in a post-flash period). Post hoc comparisons were followed up with two-sample t-tests. Either simple or two-sample t-tests were also performed to compare different conditions when the ANOVA was not necessary; alpha levels were corrected following Bonferroni methods. We performed all of our statistical tests using standard Matlab (MathWorks, Inc.) functions.

## Results

Smooth pursuit initiation is thought to involve an “open-loop” interval, of ∼100 ms, during which eye velocity is presumably not yet under visual feedback error control. However, visual signals can enter into the processing hierarchy of the oculomotor system significantly earlier than 100 ms. For example, transient visual responses in the brainstem can occur with latencies shorter than ∼40-50 ms (Rizzolatti et al., 1980; Boehnke and Munoz, 2008; Chen et al., 2018). We thus hypothesized that there exist mechanisms in which even “open-loop” smooth pursuit initiation can be significantly modified by visual inputs, and in a spatially-specific manner. We tested this by introducing a brief (∼8 ms) transient visual stimulus onset that effectively added a transient instantaneous eye position error signal to the smooth pursuit system (Materials and Methods; Fig. 1). We found that such a signal was highly effective in modifying smooth pursuit initiation. To demonstrate this, we first present cases in which the transient visual onset consisted of a full-screen white flash, and we then show results when spatially-precise position error signals were introduced by smaller transient flashes.

### Transient visual signals perturb smooth pursuit initiation velocity even during “open-loop” intervals

Our monkeys initiated smooth pursuit eye velocity without saccades through our use of a Rashbass step-ramp target trajectory (Experiment 1; Materials and Methods) (Rashbass, 1961). After a period of steady-state fixation, the pursuit target stepped in a direction opposite to the motion trajectory and instantaneously moved at constant speed (Fig. 1A), such that by the time at which smooth pursuit was initiated, there was no need for a catch-up saccade (de Brouwer et al., 2002). At a random time during the initiation phase of smooth pursuit, we presented a full-screen flash. Figure 2A shows an example of saccade-free smooth pursuit initiation with and without a full-screen flash from monkey M. The gray curve shows average radial eye velocity aligned on the onset of target motion across no-flash trials (i.e. control trials), and error bars denote 95% confidence intervals. As expected from the task design, eye velocity increased smoothly to approach target velocity, and the onset of smooth velocity acceleration occurred after a latency of ∼100 ms from target motion onset. However, when a full-screen flash occurred 50 ms after target motion onset (blue curve in Fig. 2A), the initiation of smooth pursuit was transiently disrupted (indicated by the arrow and label “Dip” in the figure); there was a transient reduction in eye acceleration starting ∼50 ms after flash onset, and almost coincident with the very earliest phase of smooth pursuit initiation. This effect can be better visualized in Fig. 2B; here, we subtracted, for every flash trial, the average radial eye velocity of the no-flash condition from the given trial’s radial eye velocity, and we then averaged all velocity “difference” curves (Materials and Methods). The average velocity difference curve showed a clear transient dip starting ∼50 ms after full-screen flash onset (error bars denote 95% confidence intervals). Importantly, we ensured that this data did not contain any microsaccades or catch-up saccades (Materials and Methods). Thus, early smooth pursuit initiation (so-called “open-loop” pursuit) is reliably modified by transient visual signals.

The transient change in saccade-free pursuit initiation was always time locked to flash onset. For example, Fig. 2C, D shows results similar to those from Fig. 2A, B above (and from the same monkey), except that the flash now happened 83 ms after target motion onset rather than after 50 ms. A similar transient decrease in eye acceleration during pursuit initiation was evident, but it now happened slightly later in the initiation phase of smooth pursuit. Similarly, Fig. 2E, F shows results with flash onset 117 ms after target motion onset, and Fig. 2G, H shows results with flash onset 150 ms after target motion onset. In all cases, there was a transient reduction in eye acceleration (marked by the label “Dip” in each figure panel), such that the eye velocity difference curves (Figs. 2B, D, F, H) always showed a negative-going transient ∼50 ms after flash onset. These transients are reminiscent of reductions in saccade and microsaccade frequency with short latency after full-screen visual flashes (Reingold and Stampe, 1999, 2000, 2002; Buonocore and McIntosh, 2008; Rolfs et al., 2008; Edelman and Xu, 2009; Bompas and Sumner, 2011; Hafed and Ignashchenkova, 2013; Tian et al., 2018). However, in our case, the changes to smooth pursuit initiation happened in a more “analog” fashion for smooth eye velocity as opposed to a more “digital” or “all-or-none” fashion for saccade frequency.

To demonstrate that the transient changes in smooth eye velocity were time locked to full-screen flash onset in both of our monkeys, we computed the velocity difference curves described above (e.g. Fig. 2B), but we did so for all flash times that we tested, and also after aligning the curves to flash onset as opposed to target motion onset. The left panel of Fig. 2I shows the results of this analysis for monkey A, and the right panel shows the results for monkey M. Note that for presentation purposes, the figure shows curves from every other flash time, but the results were identical for all flash times: regardless of flash time, its influence on smooth pursuit initiation appeared in each monkey ∼50 ms after flash onset. We statistically verified this observation by averaging the value of the curve for each flash trial in the interval 50-100 ms after flash onset and testing against zero. As expected, for both monkeys, there was reliable “inhibition”, or a negative-going deflection in the difference velocity signal, across flash onset times (results of the statistical tests are reported in the legend of Fig. 2).

The above results indicate that even open-loop smooth pursuit initiation is modifiable by visual signals. We confirmed that such modification is similar to what might happen during sustained, steady-state smooth pursuit, as was recently reported in humans (Kerzel et al., 2010). Specifically, in a subset of our experiments (Experiment 3; Materials and Methods), we presented the full-screen flash during sustained steady-state smooth pursuit. We picked an interval of smooth pursuit eye movements around the time of full-screen flash onset, and we removed any catch-up saccades present in such interval. Like described above, we also plotted radial eye velocity aligned on flash onset time (Fig. 3; error bars denote 95% confidence intervals). Full-screen flashes had a similar effect to their effect during smooth pursuit initiation: steady-state smooth pursuit eye velocity started being transiently reduced ∼50 ms after flash onset. Therefore, our results so far indicate that transient visual onsets reliably modify even open-loop smooth pursuit initiation, and in a manner similar to what happens during steady-state pursuit.

**Figure 3.**
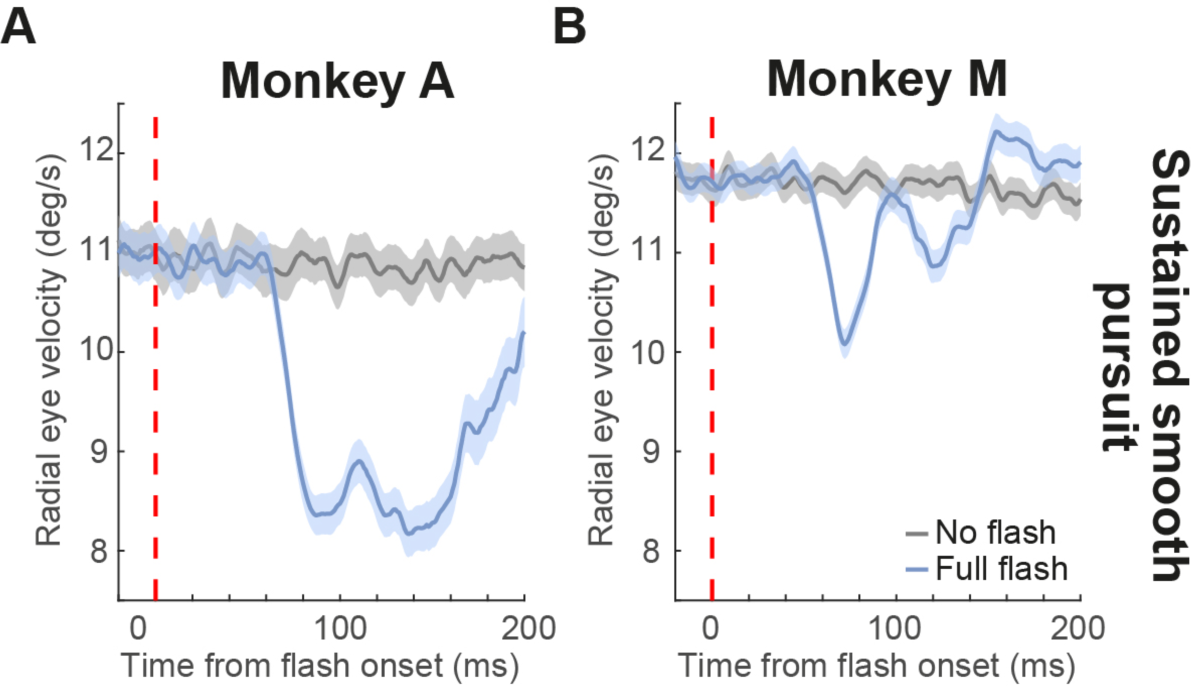
Transient disruption of sustained smooth pursuit. (A) We repeated analyses similar to Fig. 1 but during sustained smooth pursuit from Experiment 3. In monkey A, we plotted radial eye velocity with no flashes were presented (gray) or after full flash onset (light blue). There was a transient reduction on smooth eye velocity ∼50 ms after flash onset. Note that target velocity in this experiment was lower than in Fig. 2 (Materials and Methods). (B) Same for monkey M. Error bars denote 95% confidence intervals. Note that monkey A had lower sustained pursuit eye velocity than monkey M (compare gray curves in both panels). This monkey relied more on catch-up saccades. Error bars denote 95% confidence intervals. In monkey A, each velocity trace was the result of averaging 494, 548 and 530 trials for the no flash, backward flash and forward flash condition respectively. In monkey M, 892 (no flash), 974 (backward flash) and 949 (forward flash) trials.

### Transient instantaneous eye position error signals modify smooth pursuit initiation velocity in a direction-dependent manner

Given the results above, might it be possible that transient spatially-specific position error signals can also modify open-loop smooth pursuit initiation, but in a spatially-specific manner? To answer this question, we turned back to our saccade-free smooth pursuit initiation experiments of Figs. 1-2. We analyzed trials in which we presented, for only one display frame (∼8 ms), a small white square of 1 deg dimension either 2.1 deg ahead of or 2.1 deg behind the smooth pursuit target. The square could appear at different times relative to target motion onset; therefore, the flash “in front” of the pursuit target could sometimes happen either “behind” or “in front of” instantaneous gaze position depending on flash timing (Fig. 1B-D). For example, if target motion was rightward and a “forward” flash (i.e. 2.1 deg to the right of the pursuit target) happened ∼50 ms after motion onset, then the flash was still to the left of instantaneous gaze position because the target motion had not yet crossed gaze position (Fig. 1B). On the other hand, if the same flash happened ∼150 ms after motion onset, then the flash was already ahead of instantaneous gaze position (Fig. 1C). According to our hypothesis of a spatially-specific position error signal introduced by the flash, this means that both “forward” and “backward” flashes relative to pursuit target location should transiently reduce smooth pursuit acceleration if the flash happened ∼50 ms after target motion onset, but “forward” flashes should increase (rather than decrease) smooth pursuit acceleration if the flashes happened ∼150 ms after target motion onset.

This is indeed what we found. In Fig. 4A, B, we plotted data in a format similar to Fig. 2A, B, but only for “forward” flashes happening 50 ms after motion onset (from monkey M); similarly, in Fig. 4C, D, we plotted the same data for “backward” flashes happening 50 ms after motion onset (from the same monkey). In both cases, transient reductions in eye acceleration happened. On the other hand, when the flash happened 150 ms after motion onset, the transient stimulus increased eye acceleration for forward flashes (Fig. 4E, F), while it reduced it for backward flashes (Fig. 4G, H).

**Figure 4.**
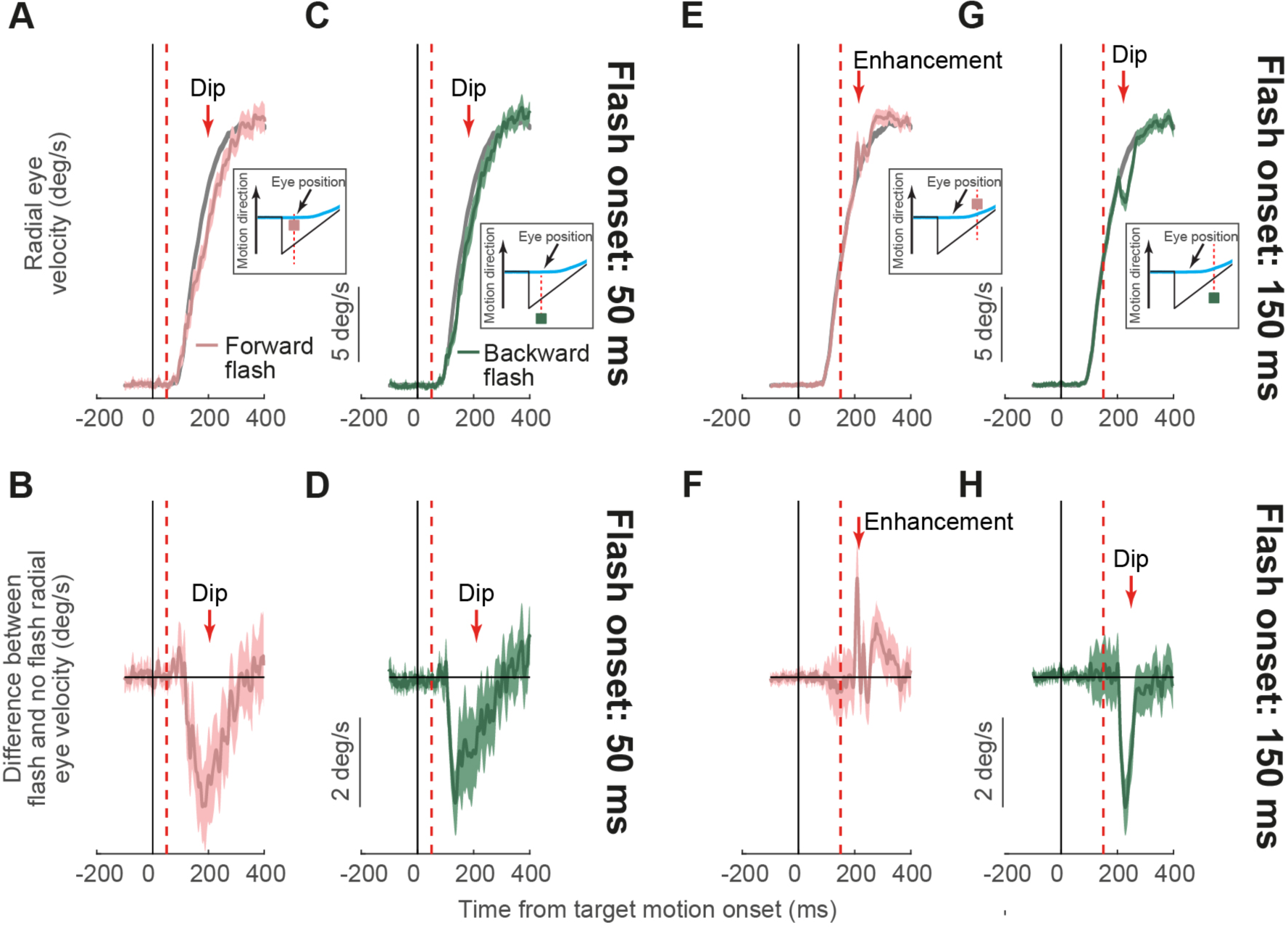
Directionally-specific modulation of open-loop smooth pursuit initiation by injected transient eye position error signals. (A, B) Example velocity (A) and velocity difference (B) traces from monkey M in the same format as in Fig. 2A, B. A flash was presented 2.1 deg ahead of the pursuit target only 50 ms after target motion onset. Because pursuit had not yet started at flash onset and because of the geometry of flash location, the flash was still behind gaze position when it occurred (see inset and Fig. 1B). This caused a transient dip as in Fig. 2A, B. (C, D) Similar analyses for a flash behind the pursuit target, again causing a transient dip as in A, B. (E, F) When the forward flash happened 150 ms after flash onset, it was now ahead of instantaneous gaze position (see inset and Fig. 1C), and now caused enhancement rather than inhibition of pursuit initiation. (G, H) A backward flash at the same time caused inhibition. Therefore, pursuit disruptions are not generalized inhibitory effects, but they reflect spatially-specific influences of flash position relative to instantaneous gaze position. Error bars denote 95% confidence intervals. After saccade removal (Materials and Methods), trial numbers for the 15 flash delays in both the forward and backward flash conditions were 7-28 (averages: 21.3 and 13.2, respectively) in monkey A, and 20-69 (averages: 52.2 and 46, respectively) in monkey M.

We summarized the observations above across both monkeys in Fig. 5A, B. For each monkey, we measured the value of the velocity difference curve between flash and no-flash trials (e.g. Fig. 2B, D, F, H and Fig. 4B, D, F, H). We did so by averaging the value of the curve for each flash trial in the interval 50-80 ms after flash onset; this gave us a population of measurements for any given flash time and flash location, and we first performed statistics by estimating 95% confidence intervals on such a population. We then plotted the velocity difference for a given flash type (e.g. forward or backward relative to pursuit target position) as a function of flash onset time relative to pursuit target motion onset (Fig. 5A, B; error bars denote 95% confidence intervals). The pink curve for each monkey shows the effect of a forward flash (relative to pursuit target position) on smooth eye velocity, and the green curve shows the effect of a backward flash. There were effects of both forward and backward flashes on even open-loop pursuit initiation. Specifically, and consistent with Fig. 4, both monkeys showed reliable influences of spatially-localized flashes on smooth pursuit initiation, with a transition from inhibition (negative y-axis values) to enhancement (positive y-axis values) in the forward flash conditions (relative to no flash pursuit initiation) according to the observations of Fig. 4. We statistically validated this result by performing a two-way ANOVA (flash time by flash location) for each of the monkeys. We found a significant interaction between the two factors of flash time and flash location (monkey A: F(14, 489) = 3.72, p = 5.902*10^−6^; monkey M: F(14, 1443) = 3.11 p = 8.499*10^-5^), supporting the hypothesis that for longer delays, the two flashes were differentiating inhibition from enhancement of eye acceleration. Starting from flashes happening at around 100 ms after motion onset (monkey A: 116 ms; monkey M: 92 ms), forward flashes led to smaller inhibition that turned into enhancement when compared to backward ones (Fig. 4). Therefore, open-loop smooth pursuit initiation is modifiable, and it reliably reflects the influence of transient instantaneous position error signals introduced by spatially localized flashes.

**Figure 5.**
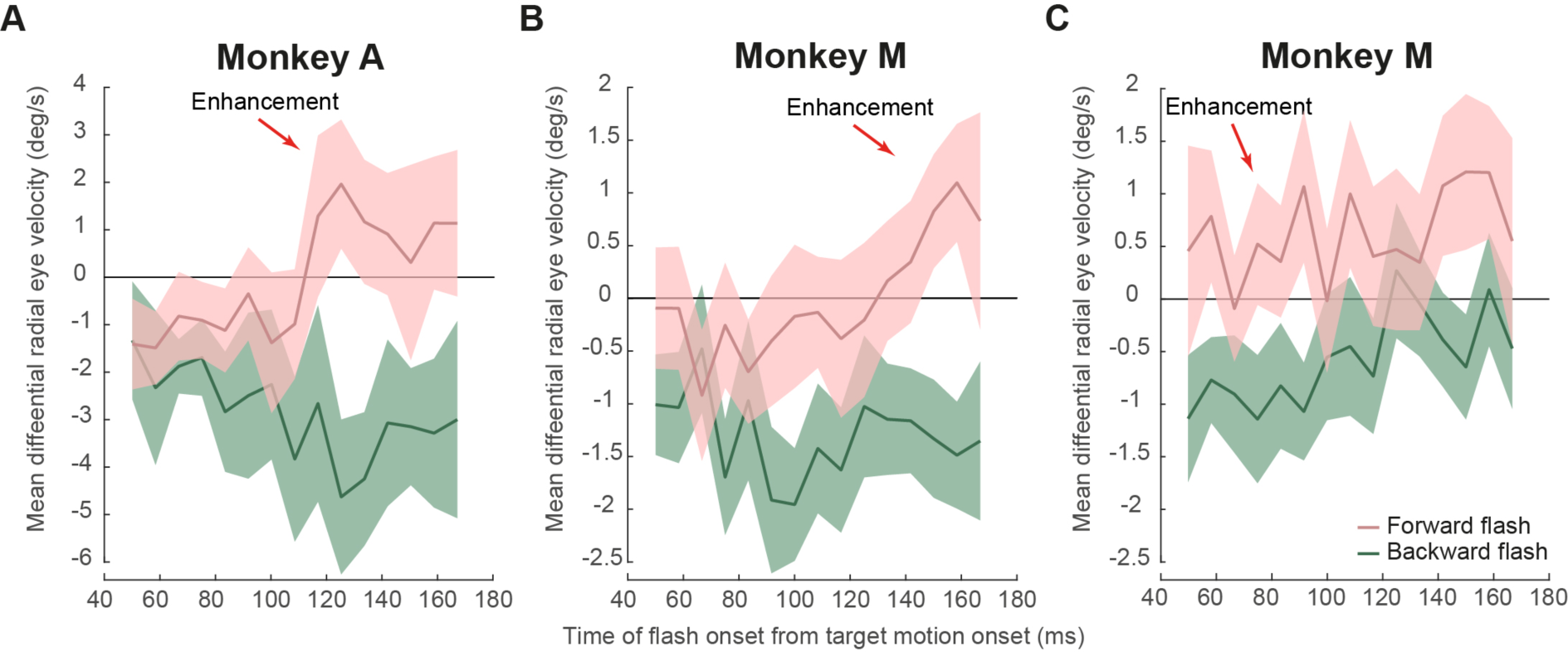
Spatially-specific transient modulations of open-loop smooth pursuit initiation depending on flash location. (A, B) For forward and backward flashes 2.1 deg ahead of or behind the pursuit target, we confirmed the observations in Fig. 4 in both monkeys. Very early flashes caused inhibition and later ones caused enhancement in smooth pursuit velocity/acceleration. For each flash time, we measured the eye velocity difference in an appropriate interval after flash onset (Materials and Methods), and we plotted this measurement as a function of flash onset time relative to target motion onset. For reference, smooth pursuit initiation typically happens at ∼100 ms, meaning that the measurements shown all illustrate modifications of open-loop smooth pursuit initiation. (C) When the flash was at 6 deg instead of 2.1 deg, enhancement occurred even for the very earliest flashes, supporting the conclusion that the early inhibition in A, B was not a generalized inhibitory effect, but that it depended on instantaneous flash location relative to instantaneous gaze location. Error bars denote 95% confidence intervals.

To show that enhancement of smooth pursuit initiation acceleration can still happen even for flashes as early as 50 ms after target motion onset, we repeated the experiment of Fig. 4 one more time, but this time, the forward or backward flashes happened 6 deg in front of or behind the pursuit target position. This means that even if the flash happened ∼50 ms after target motion onset, the flash in front of the pursuit target was also in front of instantaneous gaze position (Fig. 1D). In this case, summary plots like those shown in Fig. 5A, B showed enhancement for forward flashes even for early flashes (Fig. 5C); a two-way ANOVA (flash location by flash time) showed a predominant main effect of the factor flash location (monkey M: F(1, 1513) = 105.93, p = 4.6376*10^-24^), statistically supporting the interpretation that forward flashes induced enhancement while backward flashes led to inhibition. We also observed a main effect of flash time (monkey M: F(14, 1513) = 2.15, p = 0.0078) suggesting that later flashes had a lower inhibitory effect (in Fig. 5C, it is possible to appreciate this effect as a positive slope across time for both flash directions) and a marginal interaction between the two factors (monkey M: F(14, 1513) = 1.65, p = 0.0599).

Therefore, even the inhibitory influences seen for early forward flashes in Fig. 4A, B were not a result of a generalized inhibitory effect of flashes on eye movements, as sometimes suggested from the saccadic inhibition literature with similar flashes (Reingold and Stampe, 1999, 2000, 2002; Buonocore and McIntosh, 2008). Instead, the flash was introducing a spatially-defined geometric addition to the input of the oculomotor system, as is also the case with the saccadic system (Hafed and Ignashchenkova, 2013; Buonocore et al., 2017b). What is most intriguing is that such a spatially-specific input was able to modify even so-called open-loop smooth pursuit initiation.

Finally, for completeness, we also analyzed the effects of spatially localized flashes during sustained pursuit. Forward flashes were also effective in transiently increasing saccade-free smooth pursuit eye velocity in both monkeys, but the effect was more pronounced in monkey M (Fig. 6A, B, pink). On the other hand, backward flashes were equally effective in reducing smooth eye velocity in both animals (Fig. 6A, B, green; also see Fig. 3 for an analog of these results with full-screen flashes). These results are consistent with demonstrations that position error can modulate smooth pursuit eye velocity (Pola and Wyatt, 1980; Blohm et al., 2005), and our observations show that this happens even if “position error” is defined by multiple stimuli (the pursuit target and a more eccentric flash). Interestingly, both monkeys showed a weak, short-lived transient enhancement in smooth velocity right before the reduction after the backward flashes (Fig. 6A, B), but this enhancement effect was weaker than with forward flashes and completely absent with full-screen flashes (Fig. 3).

**Figure 6.**
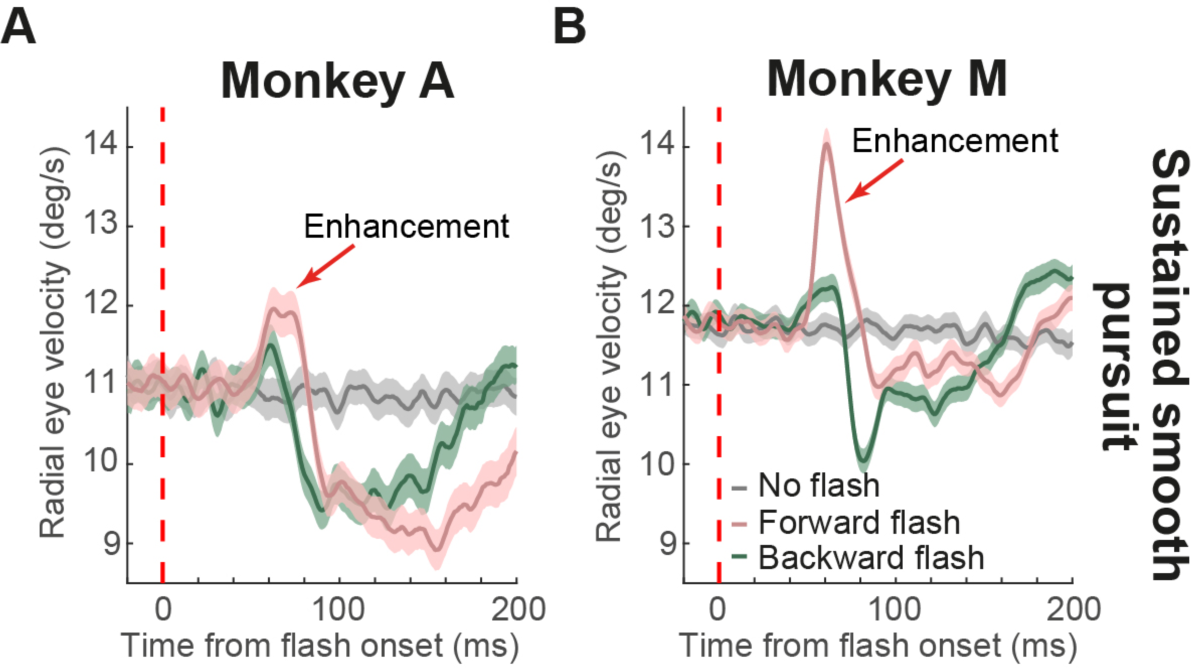
Spatially-specific modulations of sustained smooth pursuit eye movements. (A, B) In both monkeys, we repeated analyses similar to Fig. 3, but for localized flashes ahead of or behind the pursuit target (by 2.1 deg). The gray curve shows sustained pursuit velocity without flashes. The colored curves show velocity modulations with short latency after flash onset. Forward flashes caused enhanced smooth eye velocity followed by reduction. Backward flashes primarily caused only reduction in eye velocity (although there was a small, short-lived early enhancement that was not present with full flashes under similar sustained smooth pursuit conditions; Fig. 3). Error bars denote 95% confidence intervals.

In all, our results demonstrate that transient instantaneous eye position error signals can have spatially-specific effects on both open-loop (Figs. 4, 5) and sustained (Fig. 6) smooth pursuit eye movements, and that the results of Fig. 2 above are not merely a generalized inhibitory effect of flashes on eye movements.

### Catch-up saccades are influenced by transient instantaneous position error signals similarly to smooth pursuit eye velocity

We finally compared the influences of transient flashes on smooth pursuit initiation to their influences on initiation catch-up saccades. In Experiment 2, the target started moving directly from display center, resulting in the need for an initial catch-up saccade during smooth pursuit initiation. Thus, after a short latency from trial onset, there was a distribution of target-evoked saccades, whose function was to quickly catch up to the moving stimulus before smooth pursuit could proceed at full speed. We found that flashes influenced these initiation catch-up saccades similarly to how they influenced saccade-free smooth pursuit initiation. Specifically, in Fig. 7A, C, we aligned initiation catch-up saccade times to flash onset, and we separated trials based on whether the flash was in front of or behind target position. Because of the configuration of this experiment, forward flashes were ahead of instantaneous gaze position at saccade onset. We found that backward flashes caused transient inhibition of catch-up saccade frequency in both monkeys (Fig. 7A, C; dark green), and with a latency similar to that observed in smooth pursuit initiation (e.g. Fig. 4; dark green). For forward flashes, there was a brief transient increase in saccade frequency in both monkeys (Fig. 7A, C; pink, highlighted by an arrow), again similar to smooth pursuit initiation (e.g. Fig. 4; pink), before a later inhibition; full-screen flashes caused inhibition in initiation catch-up saccade frequency similar to backward flashes (data not shown). Therefore, the effects of flashes in our experiments were similar for both smooth pursuit initiation (Experiment 1) and initiation catch-up saccades (Experiment 2). The effects on saccades, in particular, were additionally reminiscent of general saccadic inhibition phenomena observed without smooth pursuit and under a variety of conditions (Reingold and Stampe, 2002; Buonocore and McIntosh, 2008; Rolfs et al., 2008; Edelman and Xu, 2009; Bompas and Sumner, 2011; Hafed, 2011; Hafed and Ignashchenkova, 2013).

**Figure 7.**
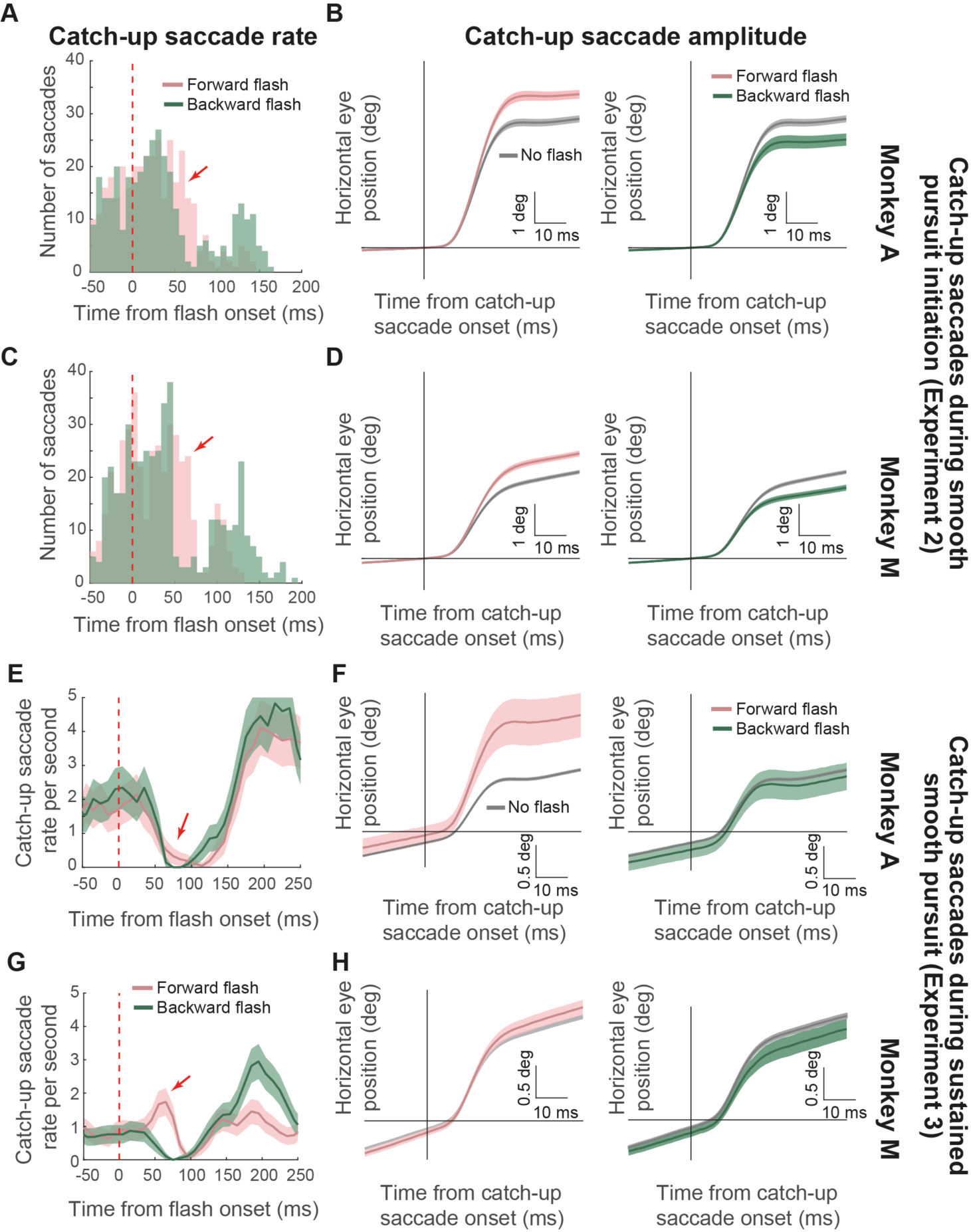
Modulations in catch-up saccades during smooth pursuit initiation and sustained smooth pursuit. (A) We plotted the distribution of initiation catch-up saccades in Experiment 2 relative to flash onset for forward (pink) and backward (dark green) flashes. Because we had similar numbers of trials across flash locations, we plotted raw, rather than normalized, histograms showing absolute trial numbers per bin (bin width: 7 ms). Backward flashes caused characteristic saccadic inhibition. Forward flashes caused an enhancement followed by inhibition. (B, left panel) We aligned eye position on initiation catch-up saccades in the same monkey and experiment. For post-flash saccades (pink), initiation catch-up saccade amplitude was larger than with no flashes (gray). (B, right panel) Post-flash initiation catch-up saccades were smaller in amplitude than without flashes for backward flashes (dark green). (C-D) Similar analyses as in A-B but for monkey M. The same observations as in monkey A were made. (E-H) The same analyses were repeated for catch-up saccades during sustained smooth pursuit. For saccade frequency (E, G), we estimated catch-up saccade rate across trials (and conditions) rather than plotting raw histograms as in A, C, because the latter histograms were counting a specific trial event (first target-evoked saccade in a trial) rather than describing an overall sustained rate that was modulated at random times by flash onsets. We replicated all of the effects in A-D, except that backward flashes during sustained pursuit (F, H, right panel) were less effective in reducing catch-up saccade amplitude than during smooth pursuit initiation. Error bars denote 95% confidence intervals.

Initiation catch-up saccade amplitudes were also modulated in a flash-dependent manner, and again consistent with our smooth pursuit initiation effects. We aligned all initiation catch-up saccades from Experiment 2 together, and we classified them as having a direction either congruent or incongruent with “forward” or “backward” flash location. For comparison, we also analyzed initiation catch-up saccades on no-flash control trials (using a virtual timing of fictive flashes that did not occur in reality). In both monkeys, initiation catch-up saccades triggered 30-80 ms after the onset of a forward flash were larger than those without a flash (two-sample t-tests, monkey A: t = 9.113; df = 255; p < 0.0001, monkey M: t = 8.411; df = 335; p < 0.0001) (Fig. 7B, D, left panels). Similarly, the saccades happening immediately after backward flashes were smaller than without a flash (two-sample t-tests, monkey A: t = −5.278; df = 217; p < 0.0001, monkey M: t = −8.205; df = 283; p < 0.0001) (Fig. 7B, D, right panels). These results are consistent with the smooth eye velocity modulations that we observed in Figs. 4, 5 during saccade-free smooth pursuit initiation. These results, for saccades in particular, are also consistent with our recent hypothesis that at the time of flash-induced visual neural activity in oculomotor circuits, like the superior colliculus (SC), not only would saccades be inhibited (e.g. Fig. 7A, C), but the few saccades that may get triggered nonetheless would also have an amplitude reflecting the addition of visually-induced neural activity to the already pre-programmed saccadic command (Hafed and Ignashchenkova, 2013; Buonocore et al., 2017b). Therefore, our smooth pursuit initiation effects were replicated in saccadic effects when tracking was initiated with the aid of initiation catch-up saccades.

We also confirmed all of the above observations during sustained smooth pursuit in Experiment 3. In this experiment, we characterized catch-up saccade frequency around the time of flash onset. As with the initiation saccades (Fig. 7A, C), there was a dip in catch-up saccade rate after flash onset in both monkeys (Fig. 7E, G). This effect had similar characteristics to previous observations in human smooth pursuit (Kerzel et al., 2010). Here, we additionally explored the influence of flash location relative to gaze position, and we observed similar effects to the initiation saccades (Fig. 7A, C) and also to the sustained smooth pursuit effects of Fig. 6. Specifically, while backward flashes strongly decreased catch-up saccade frequency in both monkeys, forward flashes caused weaker/later inhibition in monkey A and even a brief reverse effect in monkey M (increasing saccade likelihood 50-70 ms after flash onset before decreasing it). This pattern of effects mimics our smooth velocity effects, in which monkey M also showed more pronounced smooth velocity modulations after forward flashes when compared to monkey A (Fig. 6B). Catch-up saccade amplitudes during sustained smooth pursuit were also affected by the flashes in a predictable manner. We repeated the analyses of Fig. 7B, D for catch-up saccades in Experiment 3 that were triggered 50-150 ms after forward or backward flashes (amplitude dynamics in this condition were slightly slower than with initiation saccades, resulting in a slightly later analysis window). In both monkeys, forward flashes caused increased catch-up saccade amplitudes (two-sample t-tests, monkey A: t = 6.992; df = 1707; p < 0.0001, monkey M: t = 3.701; df = 1742; p = 0.0002) (Fig. 7F, H, left panels). Backward flashes, on the other hand, had a minimal impact in reducing saccade amplitude (Fig. 7F, H, right panels) (Buonocore et al., 2016; Buonocore et al., 2017b) (also see Discussion).

Taken together, our results indicate that the sudden onset of a visual transient can interfere with the spatial parameters of an ongoing motor plan (Buonocore et al., 2016; Buonocore et al., 2017b), whether saccadic or not.

## Discussion

We injected brief, perceptually subtle visual flashes around the time of smooth pursuit initiation, and we observed robust modulations in eye movements. These modulations were spatially-specific, depending on flash location relative to instantaneous gaze position, and they were therefore not reflecting a generalized inhibitory effect similar to freezing behavior in prey animals. Instead, eye acceleration during pursuit initiation was equally easy to increase or decrease, even for flashes appearing as early as ∼50 ms after target motion onset (i.e. even before the onset of smooth tracking itself). We also observed that catch-up saccades were similarly affected by flashes. Our results demonstrate that so-called “open-loop” smooth pursuit is more nuanced than generally assumed, and they motivate neurophysiological investigations of rapid sensory and motor processing in the oculomotor system.

### Beyond saccadic inhibition

Our experiments were motivated by related research investigating the impacts of brief flashes on saccades (Reingold and Stampe, 1999; Buonocore and McIntosh, 2008). Initially, such flashes were discovered to transiently inhibit saccades (Reingold and Stampe, 1999) and microsaccades (Engbert and Kliegl, 2003; Rolfs et al., 2008; Hafed et al., 2011). However, other research demonstrated that flashes also have directionally-specific influences (Hafed and Clark, 2002; Engbert and Kliegl, 2003; Hafed and Ignashchenkova, 2013; Tian et al., 2016; Tian et al., 2018), as well as effects on saccade kinematics (Buonocore et al., 2016; Buonocore et al., 2017b). For example, microsaccades occurring 50-100 ms after flash onset and directionally-congruent with its location are not inhibited as strongly as opposite microsaccades (Hafed and Ignashchenkova, 2013; Tian et al., 2016; Tian et al., 2018), and sometimes even enhanced. Moreover, congruent microsaccades are enlarged, whereas opposite ones are smaller, as if to reflect averaging of the saccade-vector command with “visual” neural activity induced by flash onset (Hafed and Ignashchenkova, 2013; Tian et al., 2016; Buonocore et al., 2017b). We observed similar effects in catch-up saccades. Our results therefore support recent suggestions that these saccades behave similarly to microsaccades (Heinen et al., 2016; Heinen et al., 2018).

Most intriguingly, even when no catch-up saccades occurred, we still observed direct correlates of the saccadic phenomena above on smooth eye movements. Smooth pursuit initiation was enhanced or inhibited depending on flash location relative to eye position, and sustained pursuit was also affected. The most interesting added advantage of performing the experiments on smooth pursuit rather than saccades was that we could focus on so-called “open-loop” saccade-free pursuit initiation, demonstrating that it is equally malleable as sustained pursuit. We specifically found that open-loop pursuit is modifiable by visual signals injecting an instantaneous transient eye position error. This observation is consistent with the original work of Lisberger and Westbrook (1985): careful reading of that study reveals that a change in target velocity led to an inflection in initial pursuit eye velocity ∼100 ms after the change. Such alteration in eye velocity also occurred after introducing position/velocity errors by means of retinal image stabilization (Morris and Lisberger, 1987). In our experiments, we extended these important findings by emphasizing the malleability of the smooth pursuit initiation phase, as well as the saccadic system, to transient visual inputs different from the target itself, even as early as 50 ms after flashes. All of this means that the term “saccadic inhibition” does not encompass the entire space of impacts of brief flashes on the oculomotor system.

In our experiments, unlike in smooth pursuit, the amplitudes of catch-up saccades after backward flashes were not always as strongly modulated as after forward flashes (Fig. 7). This may reflect the fact that smooth pursuit was ongoing. For example, backward flashes reduced smooth eye velocity itself in addition to influencing saccades (Figs. 3, 6). Therefore, there was a conflict between the increased eye position error due to slower eye velocity (supporting a bigger catch-up saccade) and the backward visually-induced neural activity of the flash (supporting a smaller catch-up saccade). Given the importance of eye position error in smooth pursuit (de Brouwer et al., 2002), it could be the case that the reduction in smooth eye velocity acted to compensate for the effects of the flash on catch-up saccade amplitude. Indeed, a plan to make a corrective eye movement coinciding with a congruent peripheral visual flash in the same direction is known in microsaccades to have maximal effects on the movement being planned (Tian et al., 2016; Tian et al., 2018).

### Smooth pursuit mechanisms

We also observed additional smooth velocity effects that we think reflect the constant interplay between target position/motion and instantaneous ongoing eye behavior. For example, smooth velocity after forward flash onset in Experiments 1 and 3 exhibited a robust post-enhancement suppression (e.g. Fig. 6). Such a decrease in eye velocity could reflect, in part, the fact that the flash-induced enhancement may cause gaze to momentarily overpass the pursuit target, resulting in negative eye position error. A more likely explanation, however, is that the deceleration might instead reflect a switch between smooth pursuit behavior and saccades. Specifically, enhancement (Fig. 6) takes place in a time window during which catch-up saccades are mostly inhibited as a result of flash onset (Fig. 7). After such inhibition, between 150 to 250 ms after flash onset, and coinciding with the smooth velocity deceleration, there is a significant rebound in catch-up saccade frequency. Thus, deceleration in pursuit velocity after the enhancement might represent a trade-off between smooth pursuit eye movements and saccades.

Our smooth velocity effects also relate to studies of distractor effects on smooth pursuit (Spering et al., 2006). We think that we complement these observations and highlight the relevance of position error for influencing smooth eye velocity. Such an influence of position error even happens during fixation with smooth ocular drifts (Skinner et al.; Martins et al., 1985; Kowler and Blaser, 1995).

In all, our results motivate interesting neurophysiology experiments that not only focus on the specification of target motion signals for smooth pursuit, but that also investigate, at the level of the brainstem, cerebellum, and cortical areas, the aggregate interplay between visual and motor signals, with and without transient flashes. One prediction out of our work is that SC recordings should reveal simultaneity of flash-induced visual activity with alterations in smooth eye velocity, but it would be interesting to know how other areas down-or upstream of SC contribute. Specifically, flash-induced visual activity is expected to appear, almost simultaneously, in many different areas whose functional specializations may support either enhancing or suppressing smooth eye velocity. What is it, then, about certain conditions (e.g. forward flashes) that make enhancement win over suppression in such a competitive scenario from the perspective of neural circuits? One such competitive scenario, which motivates our own neurophysiology experiments, relates to pre-motor brainstem omnipause neurons (OPN’s), which are believed to inhibit eye movements (Keller, 1974; Missal and Keller, 2002). During smooth pursuit initiation, OPN’s exhibit short-latency visual activity (Missal and Keller, 2002) associated with target motion onset (i.e. the visual image of the pursuit target moves near the fovea). Moreover, while this “visual” activity is not characterized in the literature, it does appear to have similar latency as SC visual activity (Evinger and Kaneko, 1982; Busettini, 2003). If we were to now record OPN activity during our tasks, we would then expect flash-induced OPN activity (inhibiting eye movements) to compete, almost simultaneously, with similar flash-induced SC activity (geometrically adding to eye movements in a manner that can enhance them). This concomitance of visual activity in different brain areas that effectively compete with each other for influencing behavior is a topic that is amenable for neurophysiological investigation, and one that we are deeply interested in. Such investigation can even clarify the curious observation that even backward flashes in our experiments sometimes caused a small early enhancement in eye velocity before the inhibition (e.g. Fig. 6, green).

### Implications for experiments

Finally, experiencing our brief flashes in pilot human experiments revealed a huge contrast between their subjective perceptual appearance to the observer and their impacts on the oculomotor system. These brief stimuli (even full flashes) seem, at face value, to be too transient and inconsequential to affect eye movement behavior (or even perceptual processing). Yet, they do, and in a most robust manner (e.g. Fig. 2). This means that certain experimental manipulations might seem so subtle to task designers but fundamentally systematic for the brain processes being invoked by the tasks. This can influence experimental result interpretations. For example, there is interest in probing perceptual state during smooth pursuit initiation by presenting brief stimuli around the time of eye movements (Spering et al., 2006; Schütz et al., 2007; Schutz et al., 2008; Braun et al., 2017). Because our results suggest that pursuit is indeed altered by brief stimuli, even during initiation, ideal stimuli for these kinds of experiments should be ones that do not alter retinal image statistics even if smooth pursuit were to be transiently slowed down or accelerated. Otherwise, experimental results may reflect differences in retinal image statistics when pursuit is altered, rather than more interesting perceptual and cognitive factors. A second similar example is the employment of brief stimulus flashes as attentional cues during fixation and assuming that microsaccades are irrelevant, an assumption that we now know to be too simplistic (Hafed et al., 2015; Tian et al., 2016).

## Acknowledgements

We were funded by the Hertie Institute for Clinical Brain Research and by the Deutsche Forschungsgemeinschaft (DFG) (Grant: FOR1847; HA6749/2-1). We also received support from the Werner Reichardt Centre for Integrative Neuroscience (CIN). The CIN is an Excellence Cluster (EXC 307) funded by the DFG.

